# Mechano-Chemo Signaling Interactions Modulate Matrix Production by Cardiac Fibroblasts

**DOI:** 10.1101/2020.05.06.077479

**Authors:** Jesse D. Rogers, Jeffrey W. Holmes, Jeffrey J. Saucerman, William J. Richardson

**Author notes:** Corresponding Author at: Department of Bioengineering, Clemson University, 301 Rhodes Research Center, Clemson, SC 29634, USA., E-mail address (W.J. Richardson).

## Abstract

Extracellular matrix remodeling after myocardial infarction occurs in a dynamic environment in which local mechanical stresses and biochemical signaling species stimulate the accumulation of collagen-rich scar tissue. It is well-known that cardiac fibroblasts regulate post-infarction matrix turnover by secreting matrix proteins, proteases, and protease inhibitors in response to both biochemical stimuli and mechanical stretch, but how these stimuli act together to dictate cellular responses is still unclear. We developed a screen of cardiac fibroblast-secreted proteins in response to combinations of biochemical agonists and cyclic uniaxial stretch in order to elucidate the relationships between stretch, biochemical signaling, and cardiac matrix turnover. We found that stretch significantly synergized with biochemical agonists to inhibit the secretion of matrix metalloproteinases, with stretch either amplifying protease suppression by individual agonists or antagonizing agonist-driven upregulation of protease expression. Stretch also modulated fibroblast sensitivity towards biochemical agonists by either sensitizing cells towards agonists that suppress protease secretion or de-sensitizing cells towards agonists that upregulate protease secretion. These findings suggest that the mechanical environment can significantly alter fibrosis-related signaling in cardiac fibroblasts, suggesting caution when extrapolating *in vitro* data to predict effects of fibrosis-related cytokines in situations like myocardial infarction where mechanical stretch occurs.

## 1. Introduction

Myocardial infarction (MI) affects a significant proportion of the US population, with approximately 8 million adults diagnosed from 2011-2014 [1]. Although short-term outcomes have improved with the widespread adoption of percutaneous coronary intervention and reduced response times [2], maladaptive scar tissue formation in the myocardium can lead to serious post-MI complications such as cardiac rupture, infarct expansion, or heart failure. These complications are dependent on a balance between tissue formation by the synthesis of extracellular matrix (ECM) and tissue degradation by a variety of proteases including matrix metalloproteinases (MMPs). While ECM synthesis is desirable at the infarct site for maintaining the structural integrity necessary to prevent cardiac rupture, excessive matrix accumulation can reduce systolic and diastolic function by increasing wall stiffness and decreasing wall conductivity, resulting in progression to heart failure [3]. Thus, controlling local ECM turnover is an essential component in promoting healthy scar formation post-MI.

Cardiac fibroblasts mediate the fibrotic response to injury by assuming an activated phenotype and synthesizing both matrix proteins and proteases, thereby controlling the balance in matrix turnover to form collagenous scar tissue [4]. Fibroblasts are highly responsive to post-infarct biochemical cues consisting of pro-inflammatory cytokines, growth factors, and hormonal agents that stimulate fibroblasts to degrade or deposit ECM. A growing body of research has also shown that fibroblast functionality is dependent on local mechanical cues; MI-induced loss of cardiomyocytes results in increased circumferential stretch at the infarct [5,6], which stimulates fibroblast activation through several mechano-sensing mechanisms [7]. While biochemical and mechanical signaling pathways have been studied individually in cardiac fibroblasts, few studies have examined the interplay between biochemical and mechanical signaling. Previous studies have shown that cytokine-stretch combinations can alter fibroblast activation beyond either cytokine or stretch treatment alone [8,9], but several questions remain regarding the role of stretch within the context of biochemical signaling, such as the relative contribution of stretch towards fibrosis compared to biochemical agonists and the ability of stretch to regulate biochemical signaling.

In this study, we address these questions by assessing the effects of six different biochemical agonists on cardiac fibroblast behavior in different mechanical contexts. Using a multi-well cell stretching device, we subjected fibroblast-seeded fibrin gels to multiple doses of each agonist under static conditions or cyclic stretch. By measuring the secretion of 13 fibrosis-regulating proteins across all mechano-chemo combinations, we examined how stretch contributes to overall fibrotic activity when combined with each agonist as well as how stretch alters agonist-mediated protein secretion. We found that stretch contributed to overall fibrotic activity when added to individual agonists by either amplifying agonist-mediated downregulation or suppressing agonist-mediated upregulation of several matrix metalloproteinases. We further analyzed these data to identify cases in which stretch alters fibroblast sensitivity towards biochemical agonists and found that stretch sensitized fibroblasts towards agonists that downregulate protease expression and dampened the effects of agonist stimulation on protease upregulation. This systems-level study points towards mechano-chemo-interactions that influence cardiac remodeling and highlights the complexity of intracellular signaling pathways in determining fibroblast phenotypes within the post-MI environment.

## 2. Materials and Methods

### 2.1. Cardiac fibroblast isolation

All animal procedures were conducted in accordance with the Institutional Animal Care and Use Committee at Clemson University. Primary mouse cardiac fibroblasts (mCFs) were isolated and cultured as previously reported [10]. Wild-type C57BL/6 mice (male, 10-12 weeks old, ∼25 g body weight) were sacrificed, and hearts were harvested and collected in Krebs-Henseleit buffer (Sigma, St. Louis, MO). The ventricles were isolated, minced, and digested enzymatically at 37 °C with Liberase TM (Roche, Indianapolis, IN). Supernatants from six successive digestion periods were collected, centrifuged at 300 x g at 4 °C, and resuspended into Dulbecco’s Modified Eagle’s Medium (DMEM, Sigma) containing 10% fetal bovine serum (FBS, Atlanta Biologicals, Flowery Branch, GA), 100 U/mL penicillin G, 100 μg/mL streptomycin, and 1 ng/mL amphotericin B (all Sigma). Cells were plated in culture flasks and incubated at 37 °C and 5% CO_2_ for 4 h. Fresh culture medium was added to remove non-adherent cells, and culture medium was replaced every 48 h thereafter.

### 2.2. Fibroblast-seeded fibrin gels

mCFs were seeded into fibrin gels using a previously reported procedure [12] and loaded gels into silicone 16-well plates (CellScale, Waterloo, ON). Briefly, mCFs were removed from flasks with 0.25% Trypsin-EDTA (Gibco, Grand Island, NY) and incubated with DiI (Invitrogen, Grand Island, NY) to visualize cell membranes after treatment. Cells were resuspended in a thrombin solution (0.44 U/mL bovine thrombin in DMEM with 0.5% FBS) and combined with a fibrinogen solution (6.6 mg/mL in 20 mM HEPES-buffered saline) at a 1:1 ratio to create a fibrin gel. Prior to plating gels, well plates were modified to contain six 1 mm x 1 mm polydimethylsiloxane posts each (PDMS, Dow Corning, Midland, MI), which have been shown to facilitate gel remodeling by fibroblasts [11–14]. Wells were additionally coated with 3.5% Pluronic F-127 (Sigma) in phosphate buffered saline (PBS, Sigma) to prevent adhesion to the well surfaces. After coating, 200 uL of fibrin gel was added to each well, and gels were incubated at 37 °C for 30 min to allow for polymerization. DMEM containing 0.5% FBS, 50 ug/mL L-ascorbic acid (Sigma), and 33.3 ug/mL aprotinin (Roche) was then added to each well, and gels were cultured for 48 h to allow for gel remodeling. For each replicate, 14 gels were seeded with 1×10^5^ mCFs for compaction, viability, and immunocytochemistry analyses, and 14 gels were seeded with 2×10^5^ mCFs to collect sufficient protein for secretomic analysis.

### 2.3. Cyclic stretch and cytokine treatment

Prepared well plates were loaded into MechanoCulture FX2 uniaxial stretching devices (CellScale, Waterloo, ON). For stretched gels, a strain amplitude of 6% and a frequency of 1.0 Hz were used to match previous cyclic stretch studies [15–17]. Axial and transverse strains for gels were validated by measuring gel deformation during 0.33-mm increments of stretch by the device actuator and calculating engineering strains from each set of values. Gels within each stretch condition were treated with either no agonist (control group) or one of six cytokines or growth factors: transforming growth factor-β1 (TGF-β1, Cell Signaling Technology, Danvers, MA), angiotensin II (AngII, Sigma), interleukin-1β (IL-1β, Cell Signaling Technology), platelet-derived growth factor-BB (PDGF, Cell Signaling Technology), L-norepinephrine bitartrate (NE, Sigma), and tumor necrosis factor-α (TNFα, Cell Signaling Technology). All concentrations were chosen from previous literature measuring fibroblast responses to individual agonists [17–22] as reported in Fig. 1C. All gels were treated for 96 h, during which doses and medium were replenished after 48 h.

**Figure 1.**
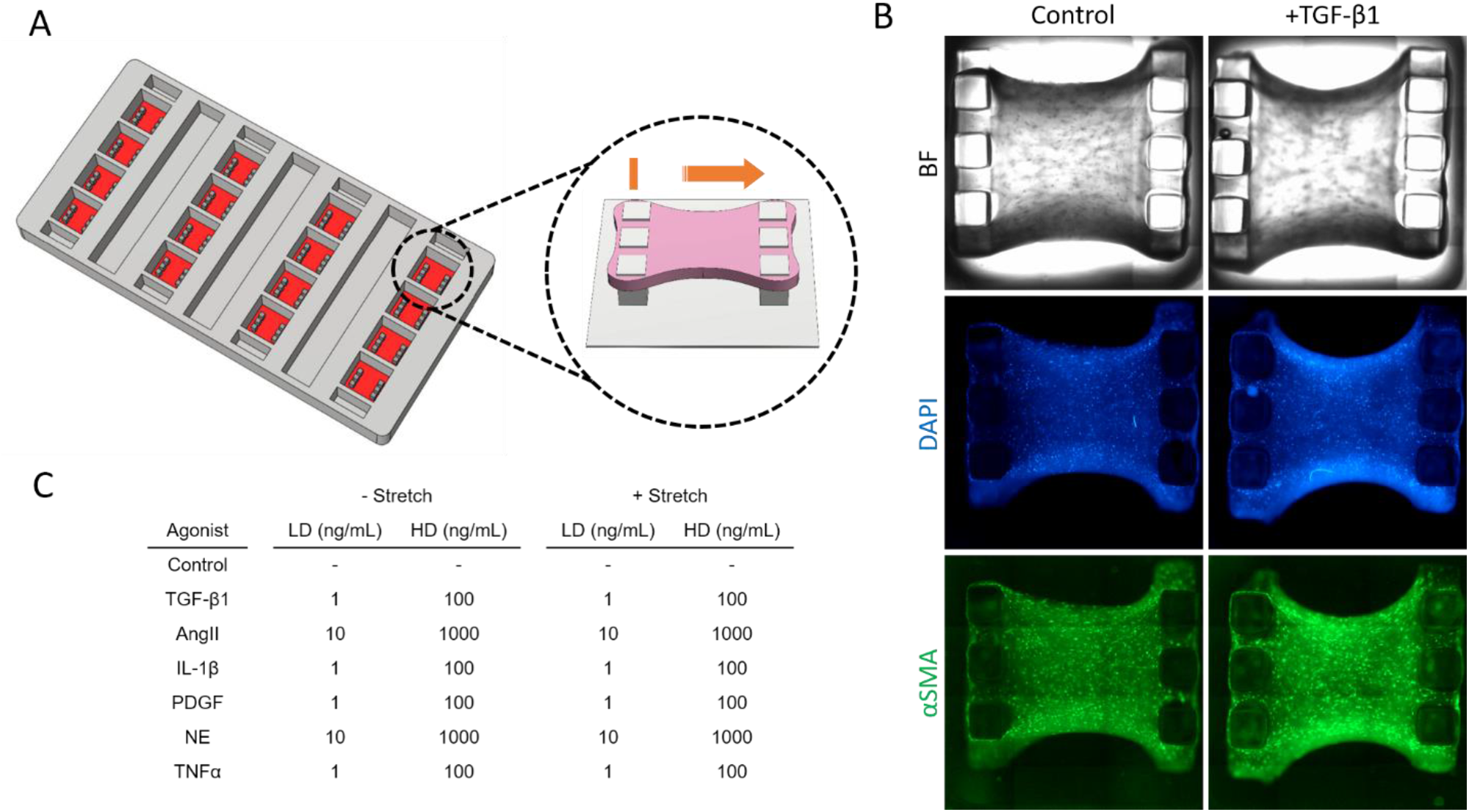
Generation and treatment of cardiac fibroblast-seeded constructs. (A) Schematic of multi-well membrane modified with silicone posts to facilitate gel remodeling and uniaxial stretch. (B) Brightfield (BF) and immunofluorescent staining for DAPI and αSMA for fibroblast-seeded gels after treatment. (C) Combinations of low dose (LD) and high dose (HD) biochemical agonists used in combination with cyclic stretch on fibroblast-seeded constructs.

### 2.4. Gel compaction analysis

Brightfield images were taken at the beginning and end of treatment using an EVOS FL Auto microscope with a 4x Plan Fluor 0.13 NA objective (Life Technologies, Grand Island, NY). A 4×3 grid of images were captured and combined to create a composite image of each gel. Gel outlines were then segmented using the threshold feature in ImageJ [23]. Gel compaction was calculated as the fold change of the final gel area from the initial gel area.

### 2.5. Immunocytochemistry and image analysis

Cell viability was determined via a fluorescence-based assay as described previously [24]. Gels were stained using a live/dead cell double staining kit (Sigma) according to the manufacturer instructions, and fluorescent images were captured at the center of each gel for calcein and propidium iodide to identify viable and dead cells, respectively. CellProfiler software was used for quantitative image analysis [25]. Viable cells were identified from calcein images using a minimum cross entropy thresholding method, and dead cells were identified from propidium iodide images using a manual threshold to remove false positives due to low numbers of dead cells per image. Cell viability for each gel was determined as the ratio of viable cells to total cells, and cell orientation was measured for viable cells as the angle from the axis of stretch to the major axis of each cell.

For analysis of fibroblast activation, immunostaining and quantitative analysis were performed as reported previously [26]. Gels were fixed in 10% neutral buffered formalin (VWR, Radnor, PA) and washed 3x in PBS. Gels were permeabilized in 0.2% Triton-X-100 (Sigma) in PBS and washed 3x in PBS, and gels were blocked with 2% bovine serum albumin (Sigma) at 4 °C overnight. Gels were stained with alpha smooth muscle actin-Alexa Fluor 488 (αSMA, Invitrogen) and 4’,6-diamidino-2-phenylindole, dihydrochloride (DAPI, Invitrogen), washing gels 3x with PBS after applying each stain. Fluorescent images were captured at the center of each gel for DAPI, αSMA, and DiI stains. Using CellProfiler software, individual cells were identified by DAPI and DiI stains, and integrated αSMA intensity was measured within each identified cell. Mean αSMA expression levels were calculated as the average integrated intensities across all cells per image. Total cell counts for each gel were extrapolated using final gel areas calculated in compaction analysis (Fig. S4A). From the number of identified cells in each image, total cell counts for each gel were calculated as the product of the cell density in each image and the final gel area.

### 2.6. Secreted protein measurement

During cell treatments, culture supernatants were collected at both 48 h and 96 h after initial treatment and stored at -20 °C. Pro-collagen I amino-terminal propeptide (PINP) and matrix metalloproteinase-1 (MMP1) content were both measured via enzyme-linked immunosorbent assay as per the manufacturer’s instructions (MyBiosource, San Diego, CA). Luminex magnetic bead assays (R&D Systems, Minneapolis, MN) were used to measure a panel of 11 additional analytes: MMP2, 3, 8, 9, and 12, tissue inhibitors of matrix metalloproteinases-1 and 4 (TIMP1 and 4), osteopontin (OPN), periostin (POSTN), thrombospondin-4 (THBS4), and plasminogen activator inhibitor-1 (PAI1). Luminex assays were measured using a MAGPIX magnetic bead reader (Luminex, Austin, TX). Absolute protein concentrations were determined from standard curves for each analyte, and all concentrations were normalized by total cell counts as determined above.

### 2.7. Agonist-stretch effect analysis

For each measured protein, contrasts between two sets of experimental groups were assessed: (i) agonist treatments alone and controls (ΔT_0_), and (ii) combined agonist-stretch treatments and controls (ΔTS). Contrasts for both sets were expressed as log_2_(fold change), and effect magnitudes were calculated as change between contrasts: *E*_*m*_ = |Δ*TS* − Δ*T*_0_|. Effect directions were calculated as the difference between contrast magnitudes: *E*_*d*_ = |Δ*TS*| − |Δ*T*_0_|. Signs of effect direction terms were used to determine categories of combinatorial effects: a positive term denoted an additive effect of combined treatment (i.e. stretch further increased or decreased protein secretion beyond agonist treatment alone), and a negative term denoted a subtractive effect (i.e. stretch suppressed the effect of agonist treatment alone). Individual contrasts that had opposite signs (e.g. ΔT_0_<0 and ΔTS>0) were denoted as inverting effects.

### 2.8. Agonist-stretch sensitivity analysis

In a similar manner to effect analysis above, contrasts between two sets of experimental groups were assessed: (i) agonist treatments alone and controls (ΔT_0_), and (ii) combined agonist-stretch treatments and samples subjected to stretch only (ΔT_s_). Contrasts for both sets were expressed as log_2_(fold change), and sensitivity magnitudes were calculated as the change between contrasts: *S*_*m*_ = |Δ*T*_*s*_ − Δ*T*_0_|. Sensitivity directions were calculated as the difference between contrast magnitudes: *S*_*d*_ = |Δ*T*_*s*_| − |Δ*T*_0_|. Signs of sensitivity direction terms were similarly used to determine categories of sensitivity changes: a positive term denoted a sensitizing effect of stretch on agonist treatment (i.e. stretch caused a greater response to agonist stimulation), and a negative term denoted a de-sensitizing effect (i.e. stretch caused a dampened response to agonist stimulation). Individual contrasts that had opposite signs (e.g. ΔT_0_<0 and ΔT_s_ >0) were denoted as reversal effects.

### 2.9. Statistical analysis

Statistical analyses were performed using the lme4 package in the R statistical suite [27]. mCFs were isolated from 8 groups of mice representing one biological replicate each, and cells were equally divided between all experimental groups within each replicate for N=8 replicates per experimental group. For each measured protein, normalized concentrations were fit to a linear mixed-effect model in which biochemical agonists and stretch states were modeled as fixed variables, and biological replicate was modeled as a random variable. Analysis of deviance was used to assess the model validity in comparison to a null model containing only random variables, using a type I error rate of 0.05 for valid models. Two-factor ANOVA was used to test fixed variables, Dunnett’s tests were used for multiple comparisons between agonists and controls, and individual Student’s t-tests with Bonferroni correction were used for comparisons made in agonist-stretch effect and sensitivity analyses.

## 3. Results

### 3.1. mCF-seeded fibrin gels secrete matrix-regulating proteins across several orders of magnitude

In order to subject fibroblasts to biochemical stimuli and mechanical stretch, we used a multi-well stretching device in which deformable culture wells transmit uniaxial tension to 3-dimensional cell-seeded fibrin constructs suspended by opposite sets of vertical posts (Fig. 1A-B). Constructs experienced an average of 6.5% axial strain and maintained viability over a 96-h treatment period (Fig. S1-2). Additionally, mCFs aligned preferably towards the direction of stretch across all experimental groups (Fig. S3), which parallels previous studies that show preferential cell orientation towards both static boundary conditions and applied strain [28,29]. Notably, TGF-β1-treated cells showed further alignment under stretch compared to static conditions, as well as increased gel compaction (Fig. S4B). Treatment with TGF-β1 has been shown to enhance myofibroblast contractility [30], which could synergize with uniaxial stretch to increase alignment although the exact mechanism remains unclear.

We used this culture platform to treat mCFs with either one of six biochemical agonists alone, cyclic stretch alone, or combinations of agonist treatments and stretch over a 96-h period to facilitate measurable protein secretion (Fig. 1C). Across all experimental groups, protein secretion varied across several orders of magnitude, from μM levels of collagen I-associated peptides to pM levels of several proteases and inhibitors (Fig. 2A). Interestingly, mCFs consistently expressed several species that have not been widely associated with fibroblast-specific protein expression, namely MMP12, TIMP4, and THBS4 [31]. Additionally, select proteins exhibit greater variance across experimental groups than others as evidenced by coefficients of variation (Fig. 2B), even within similar functional categories (e.g. MMP2/MMP9), which may be indicative of proteins with higher general sensitivity vs. robustness against stimulation.

**Figure 2.**
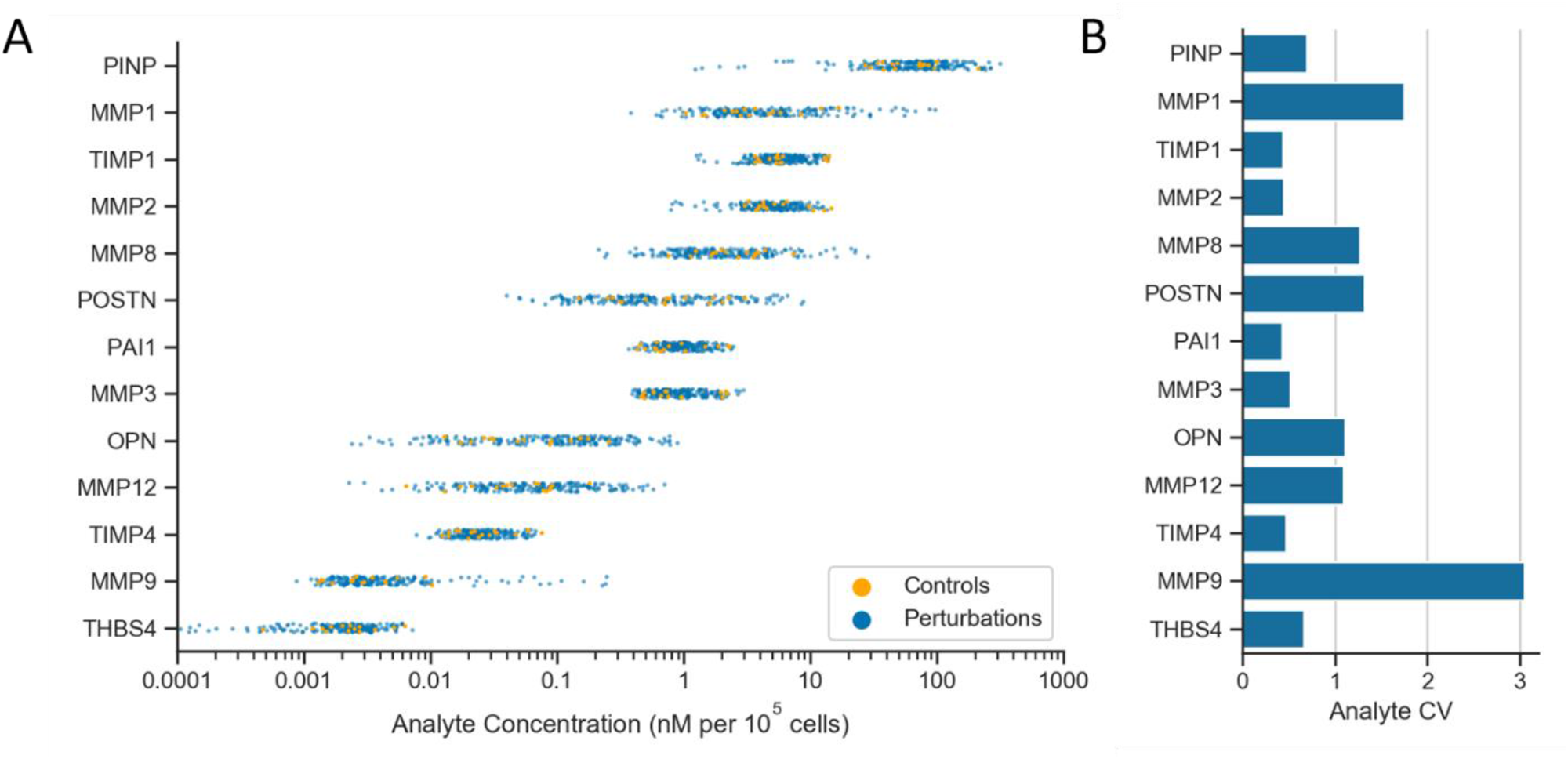
Cardiac fibroblasts express fibrosis-regulating proteins across orders of magnitude in response to biochemical and mechanical perturbations. (A) Cell count-normalized protein concentrations for un-stretched controls and all perturbations (i.e. stretched, agonist-treated, or stretched and agonist-treated). (B) Coefficients of variation (CV) for protein concentrations across all conditions.

### 3.2. Biochemical stimuli modulate fibroblast production of proteases and matricellular proteins

We examined the relationships between biochemical cues and fibrosis-related protein secretion by comparing the effects of biochemical agonists with un-stretched controls on proteins secretion. Hierarchical clustering of contrasts suggested that select proteins were highly sensitive to biochemical simulation, whereas others were more robust to these changes (Fig. 3A). Across both low and high doses, matricellular proteins POSTN and OPN, as well as MMPs 1, 8, 9 and 12, appear to be the most variably expressed. Other species were robust to changes with biochemical cues resulting in minor trends with stimulation by TNFα and PDGF, including protease inhibitors, collagen I-associated peptides and MMPs 2 and 3.

**Figure 3.**
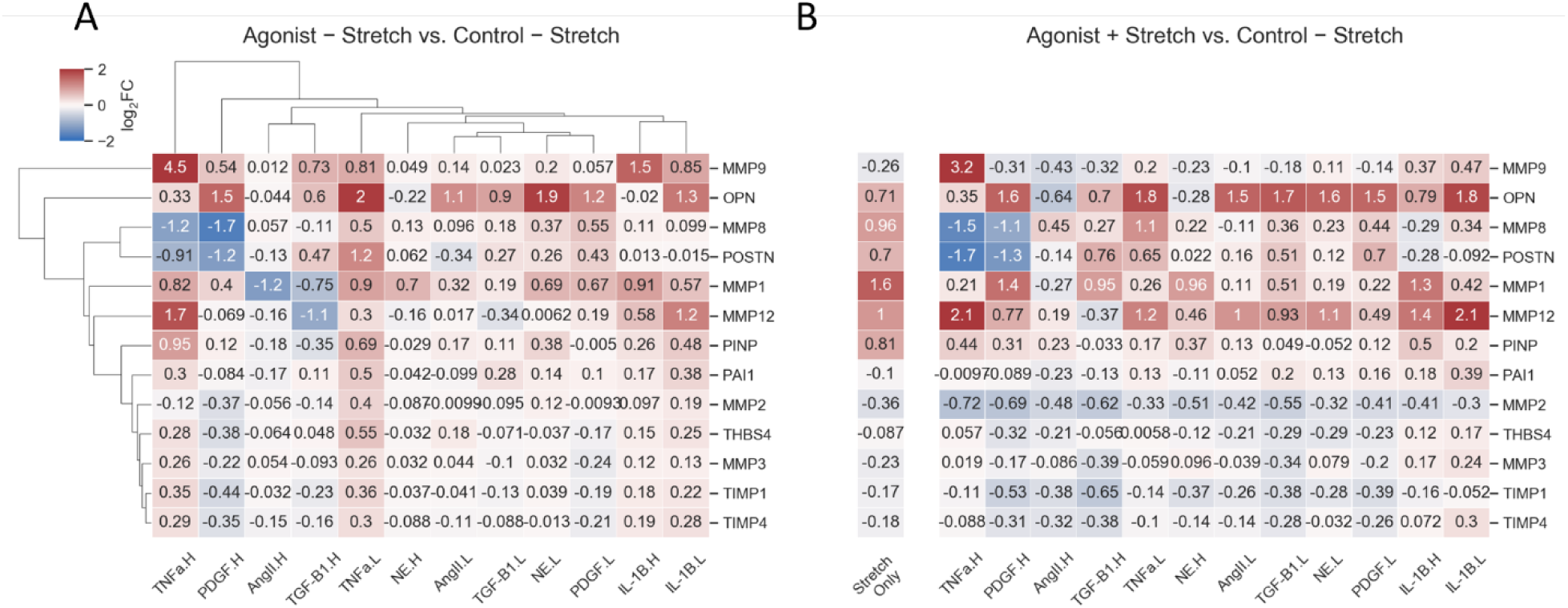
Biochemical agonists and mechano-chemo combinations alter profiles of fibroblast secretion of matrix-related proteins. (A) Changes in secretion with individual agonists. (B) Changes in secretion with cyclic stretch, either alone (left column) or in combination with agonists (right heatmap). All values are expressed as log_2_(fold-change) compared to un-stretched controls, and agonist doses are denoted by L (low) and H (high).

Across both sets of treatment doses, inflammatory cytokines IL-1β and TNFα uniquely regulated protein secretion compared to other agonist types by increasing protease production. Both agonists significantly upregulated secreted MMP9 levels (p-values shown in Fig. S5), and although levels of collagen I-associated peptides also increased with these treatments, similar increases in both MMP1 and MMP12 levels indicate that mCFs shift in balance towards the secretion of anti-fibrotic proteins. TNFα exhibited further evidence of this shift with modest increases in MMPs 2, 3, and THBS4, indicating that mCFs may be especially sensitive to this agonist.

### 3.3. Cyclic stretch synergizes with biochemical stimuli to dictate overall protein secretion

Although the importance of mechanotransduction in cardiac fibrosis has gained attention recently, it is still unclear how this signaling axis integrates with complex biochemical signaling during cardiac remodeling. We first examined cyclic stretch as an individual stimulus compared to un-stretched controls and found that stretched mCFs trended towards pro-fibrotic protein expression via upregulation of POSTN, OPN, and collagen I-associated peptides (Fig. 3B, left column). This behavior was balanced by MMP1, 8 and 12 upregulation, indicating that stretch alone did not produce definitively pro-fibrotic behavior. We then examined the role of stretch in conjunction with biochemical signaling by comparing secreted protein levels for cultures subjected to agonist-stretch combinations to un-stretched controls. mCFs exhibited similar agonist-dependent trends under stretched conditions as static cultures; matricellular proteins and MMPs 1, 8, 9 and 12 were the most variably secreted across agonists (Fig. 3B, right heatmap). Notably, inflammatory cytokines showed weaker upregulation of MMPs 1 and 9 under stretch compared to static conditions.

In order to understand how mechanical stretch contributes to overall fibroblast-mediated remodeling, we quantified differences between the effects of agonist treatments alone (i.e. agonist-only treatments vs. un-stretched controls) and agonist-stretch combinations (i.e. combinations vs. un-stretched controls). We categorized these differences in overall protein secretion into three groups: (1) stretch can have an *additive effect* by enhancing secretion levels beyond those of agonist stimulation alone (Fig. 4A, with example in 4D), (2) stretch can have a *subtractive effect* by suppressing changes in secretion with agonist treatment (Fig. 4B, with example in 4E), and (3) stretch can have an *inverting effect* by overriding the effect of agonist stimulation alone, thus changing the overall effect from upregulation to downregulation or vice versa (Fig. 4C, with example in 4F). From this analysis, we found that stretch showed trending or significant additive effects on agonist stimulation for 11 agonist-secreted protein pairs and showed trending or significant antagonistic effects for 8 pairs.

**Figure 4.**
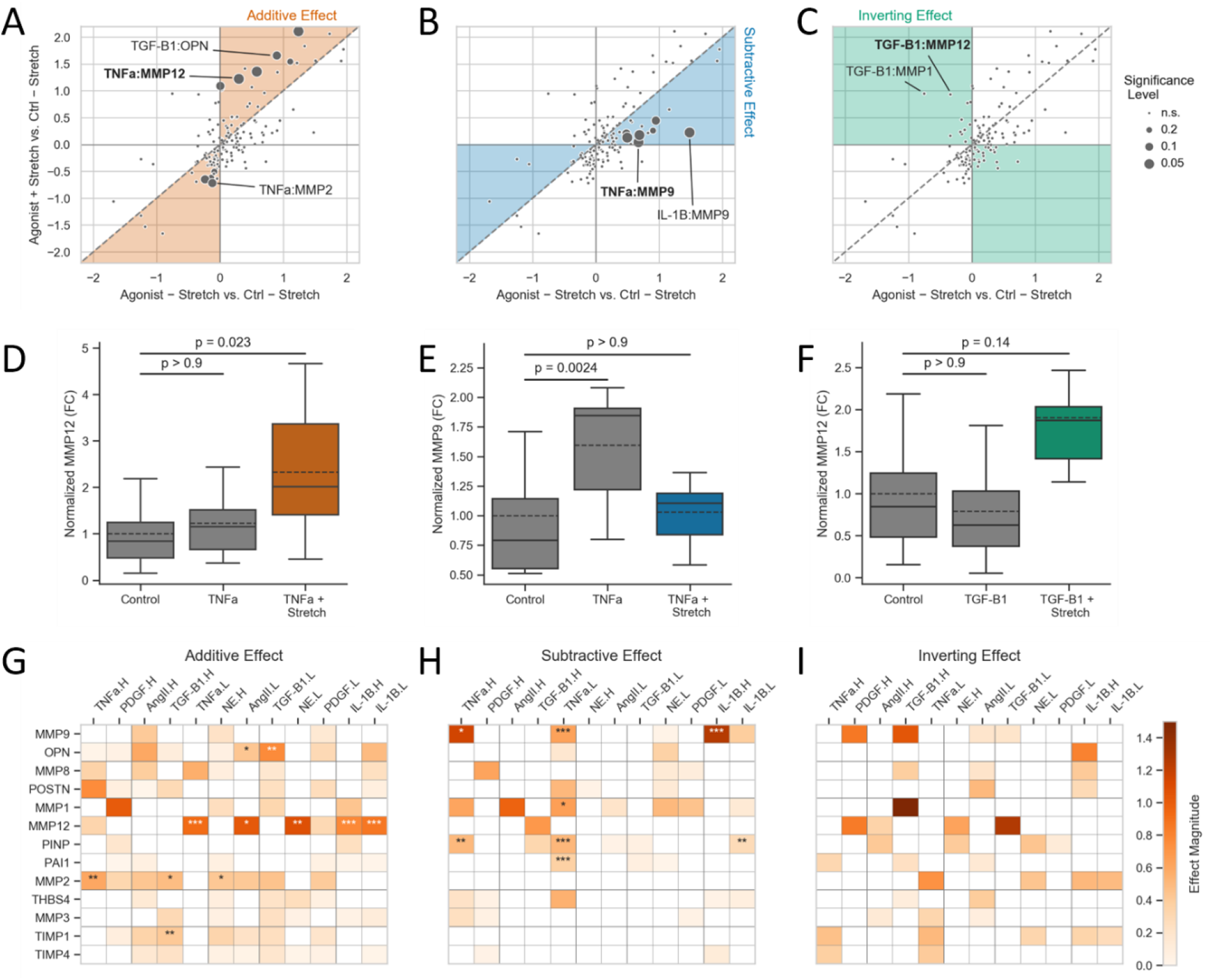
Cyclic tension differentially alters the effects of biochemical stimulation on fibroblast protein secretion. (A-C) Effects of agonist-stretch combinations on overall protein secretion. X-coordinates represent the effect of agonist treatments alone relative to un-stretched controls, and y-coordinates represent the effect of agonist-stretch combinations relative to un-stretched controls. Shaded areas indicate secreted proteins that show an additive effect compared to agonist treatment alone (A), a subtractive effect (B), or an inverting effect from downregulation to upregulation and vice versa (C). (D-F) Representative plots demonstrating effects of agonist-stretch combinations compared to agonist treatment alone. Values for all groups are normalized to mean levels of un-stretched controls. Bonferroni-corrected p-values from Student’s t-tests are used for comparisons. (G-I) Stretch effect magnitudes for cases of additive effect (G), subtractive effects (H), or inverting effects (I). Magnitudes are expressed in log_2_(fold-change), and asterisks represent level of confidence in interactions: * p< 0.2; ** p< 0.1; *** p< 0.05.

Within the additive effect group, stretch showed the most evidence of further upregulating MMP12 (Fig. 4G), with increases in expression in conjunction with IL-1β (p=0.013 for agonist/stretch combination versus un-stretched controls), TNFα (p=0.023), NE (p=0.055), and AngII stimulation (p=0.12). Secreted levels of OPN were also further increased by stretch when combined with low-dose TGF-β1 and AngII treatments, resulting in approximately 3-fold increases relative to un-stretched controls (p=0.050 and 0.11, respectively). Stretch also showed synergistic trends in a suppressive manner; when combined with TNFα, TGF-β1, and NE treatments, stretch further decreased the expression of MMP2, leading to over 2-fold reductions in protease levels compared to un-stretched controls (p=0.10, 0.16, and 0.18, respectively). By quantifying differences between agonist-only effects and combination effects across all agonist-secreted protein pairs, we found that the addition of stretch exerts the largest changes in expression for MMP12, with multiple agonist/stretch combinations increasing secreted levels approximately 2-fold above agonist-only counterparts (Fig. 4G).

Stretch demonstrated subtractive effects on protein secretion by suppressing the secretion of matrix metalloproteinases that were upregulated by agonist treatments alone (Fig. 4B). MMP9 upregulation by several agonists was especially suppressed by stretch, as both IL-1β and TNFα significantly reduced expression in combination with stretch compared to treatment with either cytokine alone (Fig. 4H). However, stretch also demonstrated antagonism towards the secretion of collagen I-associated peptides and protease inhibitor PAI1, as upregulated secretion of these species by IL-1β or TNFα were suppressed to near-control levels. Several agonist-secreted protein pairs also trended towards an inverting effect in which agonist/stretch combinations upregulated the expression of proteins that were downregulated by agonists alone (Fig. 4C). Although not meeting our criteria for statistical significance, TGF-β1-mediated secretion of MMP12 showed the most evidence of this behavior (Fig. 4F). These cases could also prove important in determining the full effect of stretch on cardiac remodeling, and further studies are necessary to explore these trends.

### 3.4. Cyclic stretch modulates fibroblast sensitivity to biochemical stimulation

In order to quantify the ability of stretch to regulate biochemical signaling, we adapted the analysis discussed above to compare the effects of agonist treatments in a static context (i.e. agonist-only treatments vs. un-stretched controls) and in a stretched context (i.e. agonist/stretch combinations vs. stretch-only treatments). We categorized changes between contexts into three groups: (1) stretch can *sensitize* fibroblasts towards biochemical stimuli by enhancing the effect of agonist treatment under stretch (Fig. 5A, with example in 5D), (2) stretch can *desensitize* fibroblasts by suppressing the effect of agonist stimulation (Fig. 5B, with example in 5E), and (3) stretch can *reverse* fibroblast behavior completely by causing agonists to downregulate secretion compared to upregulation in a static environment, or vice versa (Fig. 5C, with example in 5F). We found that stretch showed trending or significant evidence of agonist sensitization for 12 agonist-secreted protein pairs and showed similar evidence of agonist de-sensitization for 12 pairs compared to the same treatments under static conditions. None of the reversals were statistically significant.

**Figure 5.**
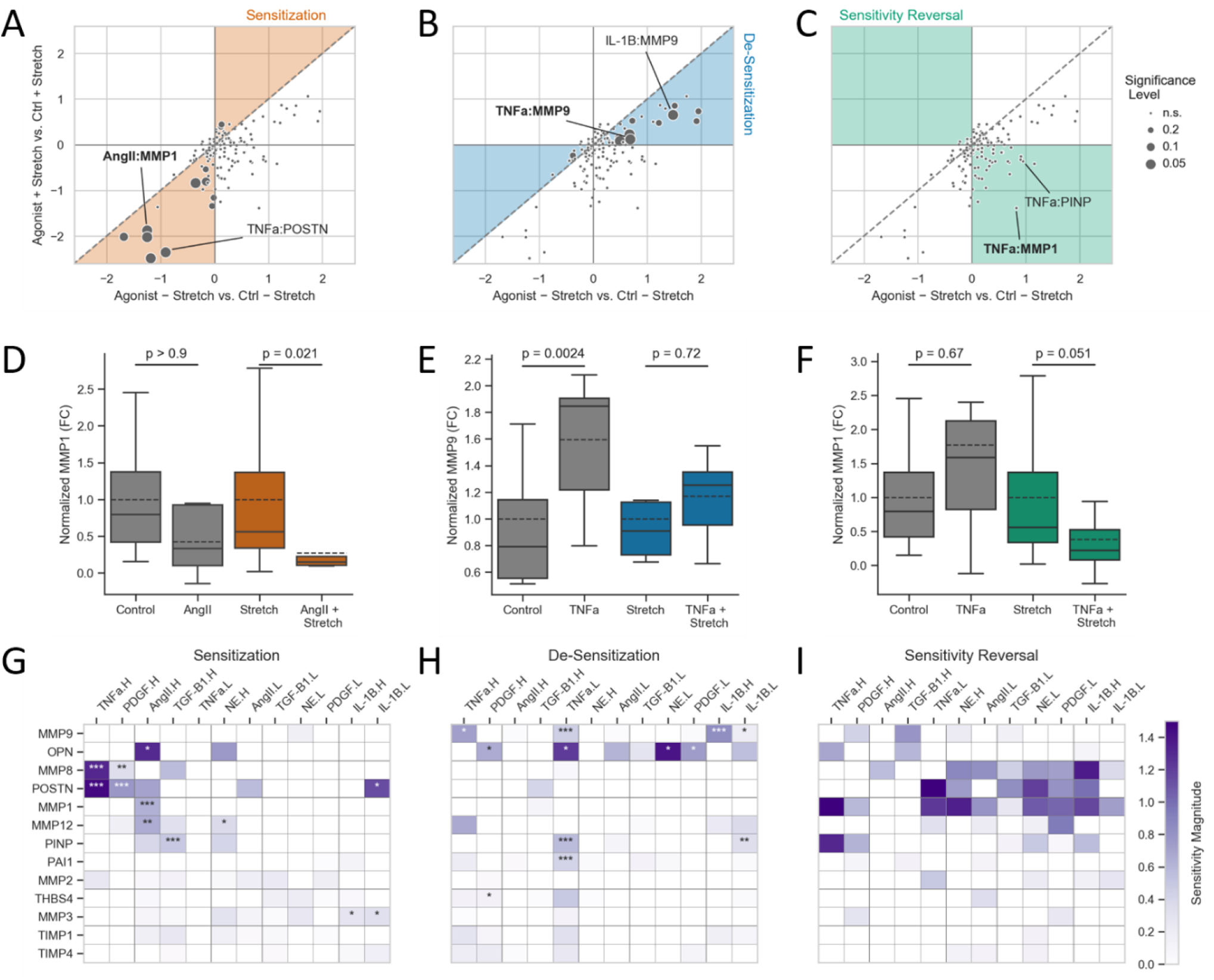
Cyclic tension can alter cardiac fibroblast sensitivity to biochemical agonists. (A-C) Changes in agonist effect on protein secretion for fibroblasts in static or strained cultures. X-coordinates represent effects of agonist treatments relative to un-stretched controls, and y-coordinates represent effects of agonist-stretch combinations relative to samples subjected to stretch only. Shaded areas indicate secreted proteins that show sensitization towards agonists (A), desensitization (B), or sensitivity reversal from upregulation to downregulation (C). (D-F) Representative plots demonstrating stretch-mediated changes in sensitivity of fibroblasts to agonist treatment. Values for un-stretched groups (gray) are normalized to mean levels of un-stretched controls, and values for stretched groups (colors) are normalized to mean levels of stretch-only groups. Bonferroni-corrected p-values from Student’s t-tests are used for all comparisons. (G-I) Sensitivity magnitudes for cases of sensitization (G), desensitization (H), or sensitivity reversal (I). Magnitudes are expressed in log_2_(fold-change), and asterisks represent level of confidence in interactions: * p< 0.2; ** p< 0.1; *** p< 0.05.

Within the sensitizing group, stretch primarily acted to amplify agonist-mediated suppression, mediating further suppression of MMP8 levels by TNFα (p=0.044 for agonist/stretch combination versus stretched-only samples) and PDGF (p=0.059). Multiple agonists also exhibited amplified suppression of POSTN secretion with stretch, with TNFα, PDGF, and IL-1β all showing trending or significant suppression in a stretched context (p=0.020, 0.034, and 0.16 respectively). Interestingly, AngII appeared to further suppress MMP1, MMP12, and OPN under stretch, with MMP1 levels decreasing to almost 25% of those for stretch-only samples (Fig. 5D). Of these trends, secreted outputs MMP8 and POSTN appeared to have the highest degrees of sensitization, as TNFα and IL-1β reduced expression of these proteins by approximately 2.5-fold more under stretch than under static conditions (Fig. 5G).

Fibroblasts also showed decreases in sensitivity towards select agonist treatments under stretch (Fig. 5B); in particular, expression of MMP9 and OPN, both of which were upregulated in response to multiple agonists in static conditions, were not significantly altered by the same agonists under stretch (Fig. 5E). Interestingly, secretion of PINP and PAI1 also showed significant desensitization towards low-dose TNFα stimulation under stretch, although magnitudes of these changes were less than those for TNFα-mediated MMP9 secretion (Fig. 5H).

We additionally found that several agonist-secreted protein pairs demonstrated a reversal in sensitivity in which agonists that upregulated protein expression in static conditions downregulated expression under stretch (Fig. 5C). Similarly to our findings of overriding effects described above, these cases did not meet our criteria for statistical significance. However, this type of behavior appears to be consistent for MMP1, MMP8, and POSTN expression (Fig. 5I), with TNFα stimulation reducing MMP1 expression to roughly 40% of stretch-only levels (Fig. 5F). Further studies validating these trends are necessary to fully understand the ability of stretch to reverse the effects of biochemical stimulation.

### 3.5. Cyclic stretch differentially regulates global fibroblast sensitivity to cytokine stimulation

We lastly assessed the sensitivity of each secreted protein to all biochemical agonists in either static or stretched conditions to identify global trends in mechano-chemo interactions. Using a summed fold-change metric for total sensitivity, we found that matricellular proteins OPN and POSTN as well as MMPs 1, 8, 9, and 12 ranked as the most sensitive to biochemical treatment in both mechanical conditions (Fig. 6A). Within these outputs, OPN, MMP9, and MMP1 all showed de-sensitization towards all agonists under stretched conditions, although total sensitivity levels still ranked highly overall. Alternatively, both POSTN and MMP8 showed increased sensitivity to agonist stimulation in a high-stretch environment, indicating that these species may be more readily regulated within infarcted areas. Using a similar analysis to compare agonist regulation across all secreted outputs, we found that stretch altered the overall influence of TNFα, NE, and AngII on the secretion of all proteins (Fig. 6B). While stretch decreased the total magnitude of effects from TNFα stimulation, however, both NE and AngII exerted greater changes in protein levels within a stretched environment, more than doubling the influence of AngII on protein secretion.

**Figure 6.**
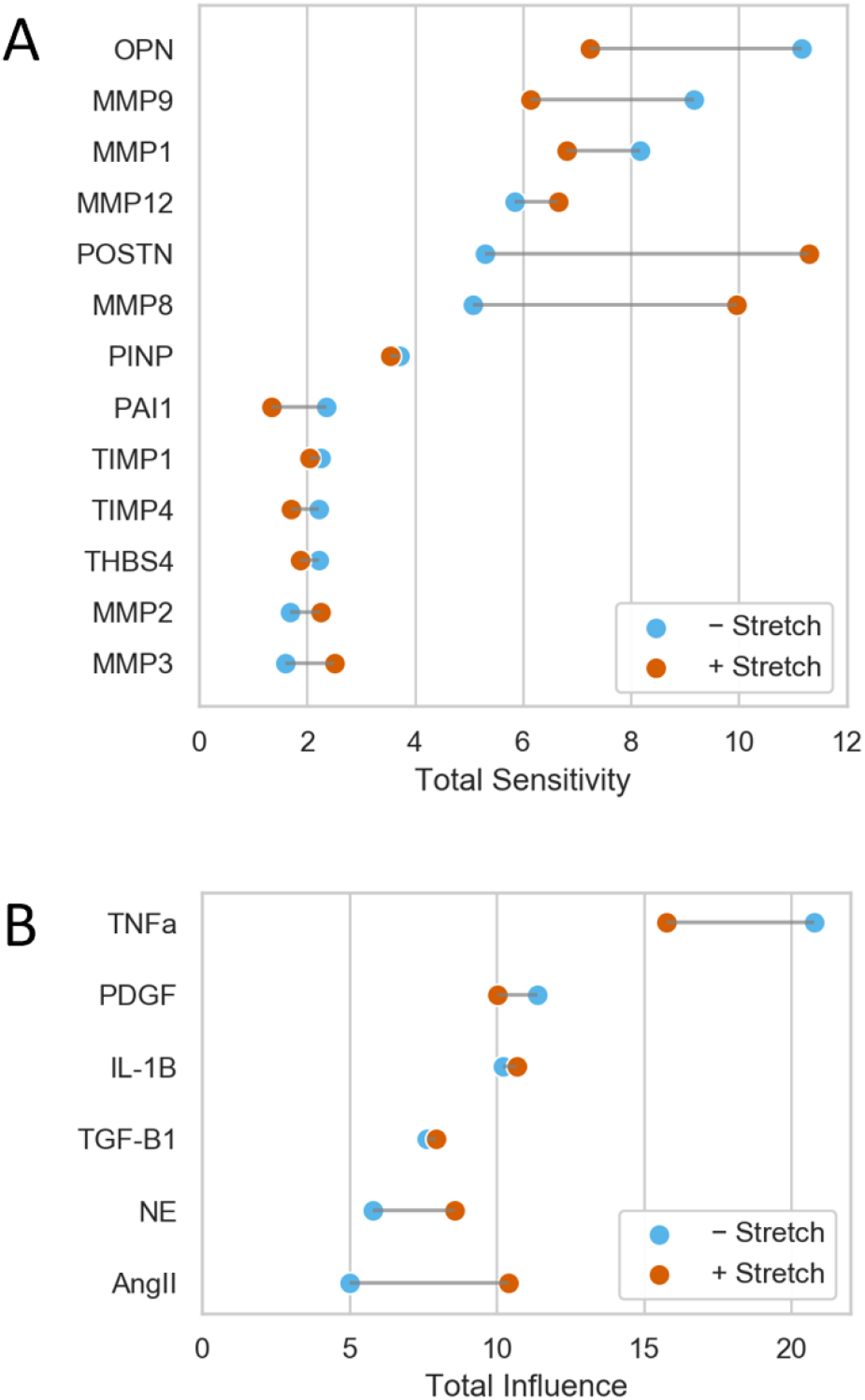
Cyclic stretch modulates overall secretion of matrix-related proteins. (A) Sensitivity of individual secreted proteins to all biochemical agonists in both stretch contexts. (B) Influence of individual agonists across all secreted proteins in both stretch contexts. Values for respective analyses are ranked by un-stretched levels for each species.

## 4. Discussion

### 4.1. Study design

The contribution of fibrosis to heart failure after MI has gained attention due to its role in chronic mechanical and electrical dysfunction. Prior studies of fibroblast behavior have highlighted the central role of dynamic mechanical stretch in the development of scar tissue, and future drug therapies that account for cell responses to this mechanical environment as well as responses to biochemical signals could improve patient outcomes. By screening the effect of individual biochemical stimuli for their effects on fibroblast-specific matrix remodeling in the presence and absence of cyclic stretch, we sought to generate new insight into stretch-mediated fibrosis within the complex post-infarct environment.

In order to subject fibroblasts to biochemical stimuli and stretch while simulating myocardial architecture and material properties, we employed a previously developed strategy in which cells are incorporated into a fibrin matrix and seeded into culture wells containing vertical, opposing sets of silicone posts. Fibroblasts cultured in this manner have been shown to remodel the surrounding matrix and form aligned constructs with comparable stiffness to myocardial levels [11,12,32,33], thus mimicking *in vivo* mechanical conditions more closely than 2-dimensional mechanical testing systems.. Although mCFs assumed an activated state throughout treatment, likely due to prior culture with stiff tissue culture flasks, the use of soft gels prevented further stiffness-related stimulation during treatment, and myofibroblast-mediated remodeling could be examined without confounding mechanical factors.

### 4.2. Relative secretion levels of matrix proteins and proteases

We first aimed to contextualize the effects of environmental cues on fibroblast-mediated scar formation by comparing absolute concentrations of mCFs-secreted proteins across and within functional categories. Collagen I-associated peptides were predominately secreted along with MMP1, MMP2, MMP8, and TIMP1, which although secreted at distinct stages after myocardial injury [34], point to dominant mechanisms of ECM turnover via fibrillar collagen synthesis collagenase-mediated degradation. We also detected relatively small, but consistent levels of MMP12, TIMP4, and THBS4, all of which have been observed in fibroblast-specific gene expression studies but have not yet been directly measured at the protein expression level [31,35–37]. These biomarkers have been implicated in post-infarct remodeling either through paracrine signaling toward resolving inflammation as in the case of MMP12 [38], or by modulating responses to pressure overload as in the case of THBS4 [39]. Their direct link to cardiac fibroblast secretion suggests that fibroblasts may act in a communicatory role in addition to directly mediating matrix turnover.

### 4.3. Differential sensitivity of secreted proteins and influence of biochemical stmuli

We compared fibroblast-specific matrix remodeling across biochemical stimuli by measuring matrix-related protein expression for six canonical agonists that have been shown to individually mediate fibroblast function [40]. MMPs 1, 8, 9, and 12 and matricellular proteins OPN and POSTN were highly sensitive to agonist treatments compared to all other measured proteins. Although absolute concentration levels were small for several proteins relative to other secreted outputs, large changes in expression suggest that select proteases may play an essential role in mediating biochemically driven matrix turnover.

Both inflammatory agonists used in our study exhibited a unique influence on protease expression relative to other agonists. We observed that both IL-1β and TNFα mediate significant increases in MMP9 levels, with TNFα additionally upregulating MMP12 expression. While these increases are accompanied by increases in collagen I-associated peptides, PAI1, and OPN, the relatively low magnitudes these changes suggest that inflammatory factors primarily acted to upregulate protease expression. These trends agree with parallel increases of inflammatory and protease levels within the acute post-infarct phase, which form the basis for removing necrotic tissue from the infarcted area [41–43].

### 4.4. Contribution of tension to overall protein secretion

Mechanical stretch has been widely shown to induce pro-fibrotic phenotypes in cardiac fibroblasts as one component of overall mechanical stimulation [44]. While previous work has shown that tensile forces can modulate fibroblast responses when combined with biochemical stimulation [8,9], it is still unclear how stretch contributes towards overall remodeling compared to biochemical signaling, and whether mechanical and biochemical signaling axes operate dependently or independently of each other. We aimed to answer these questions by treating mCF cultures with combinations of biochemical agonists and mechanical stretch and comparing the effects of these combinatory treatments to those of agonist-only treatments on protein expression.

We found that cyclic stretch altered protein secretion in two primary modalities: stretch either added to the effect of biochemical stimulation by further upregulating or suppressing protein secretion, or stretch subtracted from biochemically-mediated effects by suppressing expression levels to near-control levels. Stretch further increased expression of MMP12 and OPN compared to biochemical stimulation whereas secreted levels of MMP2 were further downregulated, suggesting that stretch reinforces pro-fibrotic activity in fibroblasts by suppressing matrix proteases and increasing matricellular protein production. This behavior was echoed by significantly suppressed MMP9 expression with stretch compared to agonist treatments alone, and although we observed similar trends for PINP levels, the relatively high magnitudes of MMP9 suppression suggest a shift in the balance of matrix turnover towards matrix formation. MMP12 levels were uniquely amplified with stretch compared to cytokine treatment alone, which could be indicative of further negative feedback by MMP12 as knockdown studies in post-MI mice have shown both an increase in inflammatory signals in the myocardium and a decrease in beneficial remodeling [38].

### 4.5. Stretch-dependent sensitivity to biochemical stimulation

We additionally tested the hypothesis that stretch can not only act alongside biochemical stimuli but also alter cellular responses to these stimuli, as previous studies have demonstrated similar signaling mechanisms in both chemo- and mechano-transduction, such as mitogen-activated protein kinase and Akt signaling pathways [32,45–47]. We found that stretch altered fibroblast sensitivity towards agonists by either sensitizing cells towards agonists that suppress protease expression (e.g. MMP1, MMP8) or by de-sensitizing cells towards agonists that upregulate expression (e.g. MMP9). These changes in protease-related sensitivity indicate that stretch interacts with chemo-transduction pathways to selectively dampen matrix protease expression. While we also observed de-sensitization of both OPN and PINP upregulation towards biochemical stimuli under stretch, stretch alone upregulated the expression of both proteins, suggesting that these ECM components remained upregulated regardless of biochemical cues in a stretched environment. These results suggest that the underlying network of fibroblast signaling plays a large role in dictating fibroblast responses to competing cues, and further studies examining the intracellular signaling network are necessary to fully understand the mechanisms of action guiding fibroblast-mediated remodeling.

Our findings suggest that cardiac fibroblasts act in a mechano-adaptive manner to regulate matrix turnover, with mechanical stretch acting to alter matrix protease and matricellular protein secretion both directly, acting independently of biochemical signaling, and indirectly, acting to modify biochemical signaling. We found that TNFα-mediated protein secretion showed the highest degree of mechano-adaptive behavior compared to the other agonists tested, with stretch-mediated expression of MMP8, MMP9, MMP12, OPN, POSTN, PINP, and PAI1. Similar findings were recently found by Aguado and colleagues in which TNFα-mediated deactivation of cardiac fibroblasts was amplified depending on substrate stiffness, with a higher proportion of cells reverting towards quiescence on stiff hydrogels [48]. Our evidence of stretch-mediated sensitization and de-sensitization towards TNFα suggests that TNFα signaling mediates fibroblast activation under dynamic mechanical conditions. Further studies are necessary to validate these relationships between inflammation and stretch in guiding fibroblast phenotypes.

### 4.6. Study limitations and future directions

One limitation of this study arises from the variability that arises from the measurement of protein expression across biological replicates. The use of separate pools of mCFs in our comparisons allowed for genomic variability between replicates, and although our use of a mixed-effect statistical model was able to discern agonist- and stretch-mediated behavior, we cannot rule out additional mechano-chemo interactions that we were unable to detect. However, the interactions that we identified in our analysis persisted despite this variability, thereby improving our confidence in these findings. Our screen of mechano-chemo interactions was additionally limited to two doses of biochemical stimulation and two mechanical states, and further dose-response studies would further explore the extent of relationships between these stimuli. Additionally, a number of other secreted factors were not measured in our study that have been shown to play significant roles in matrix remodeling, including those related to matrix processing and turnover (e.g. cathepsins, lysyl oxidase, etc.) and paracrine signaling (e.g. tenascin-c, interleukins, thrombospondins, etc.). Further measurements would provide insight into the contribution of fibroblast secretion towards maladaptive remodeling.

Lastly, our data suggest that fibroblast secretion relies on a complex network of intracellular signaling, which contains interactions between transduction pathways that dictate overall responses. Several groups have successfully identified influential mechanisms of mechano-transduction and mechano-chemo crosstalk using computational models of cell signaling [26,49,50], providing new directions towards controlling cell behavior in a dynamic mechanical environment. Future work integrating these systems-levels models with cell signaling datasets are necessary to fully elucidate mechanisms-of-action for fibroblast-mediated matrix remodeling.

## 5. Conclusions

We developed a screen of cardiac fibroblast protein secretion in response to combinations of biochemical agonists and mechanical stretch to investigate how cyclic stretch influences fibrotic activity within the context of biochemical signaling. We found that biochemical stimuli primarily altered the expression of MMPs 1, 8, 9, 12, OPN, and POSTN, and inflammatory agonists IL-1β and TNFα particularly promoted protease secretion, thereby shifting the balance in matrix turnover towards degradation. The addition of stretch to biochemical stimuli shifted fibroblast behavior towards matrix accumulation by either further upregulating matricellular protein expression, further downregulating protease expression, or antagonizing the upregulation of MMP9 by inflammatory agonists. We also found that stretch mediated fibroblast sensitivity towards biochemical stimuli by increasing sensitivity towards agonists that suppress MMP1, MMP8, and POSTN while decreasing sensitivity towards those that upregulate MMP9, thereby priming cells towards matrix accumulation. The screening approach employed here allows for the identification of emergent interactions between cell stimuli and can be used to inform targeted studies for interactions that govern matrix turnover.

## Funding

This study was supported by the National Institutes of Health (GM121342, GM103444, HL144927) and the American Heart Association (17SDG33410658).

## Disclosures

None.

## Acknowledgements

We thank Joseph Bible Ph.D. for his assistance in developing the statistical model used in this study, and the Godley-Snell Research Center for assisting in cardiac fibroblast isolation.

## Supplementary Materials

**Supplementary Figure S1.**
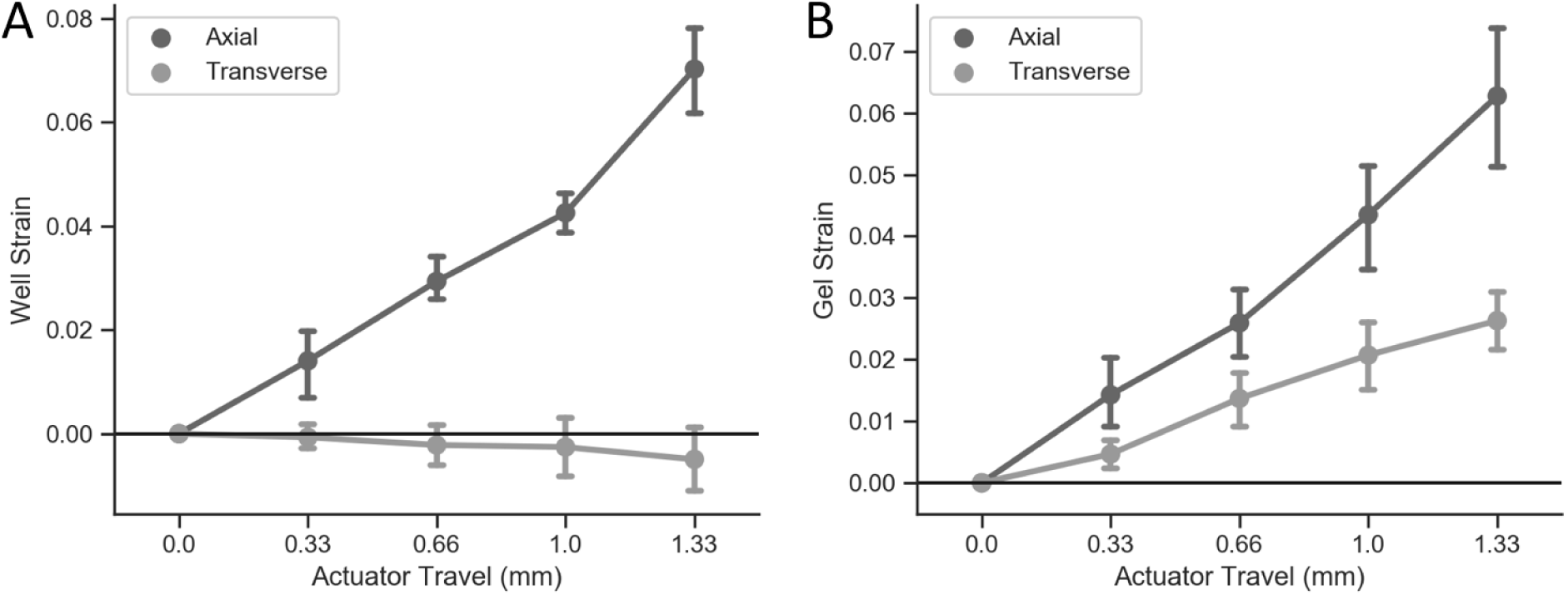
Validation of strain for PDMS wells and fibroblast-seeded gels during cyclic uniaxial tension. (A) Average strain of individual PDMS wells subjected to uniaxial tension. Engineering strains were derived from the deformation of well sidewalls in the axial and transverse directions (N=4 technical replicates). (B) Average strain of individual fibroblast-seeded fibrin gels subjected to uniaxial tension. Engineering strains were derived using similar directional conventions as in S1A (N=4 technical replicates).

**Supplementary Figure S2.**
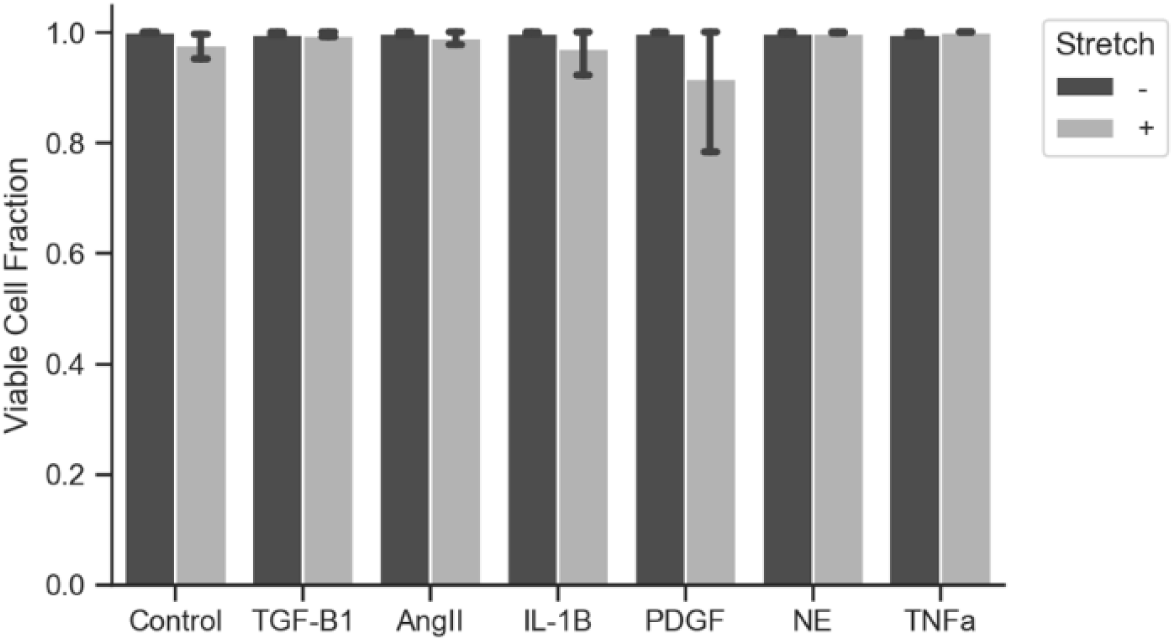
Viability of fibroblast populations across all experimental groups and biological replicates. Viability was determined using a live/dead fluorescent double staining kit after the duration of each treatment regimen and is shown as the proportion of calcein-positive cells to total cells for images taken of each gel (400.2±183.9 cells per image, N=7 biological replicates).

**Supplementary Figure S3.**
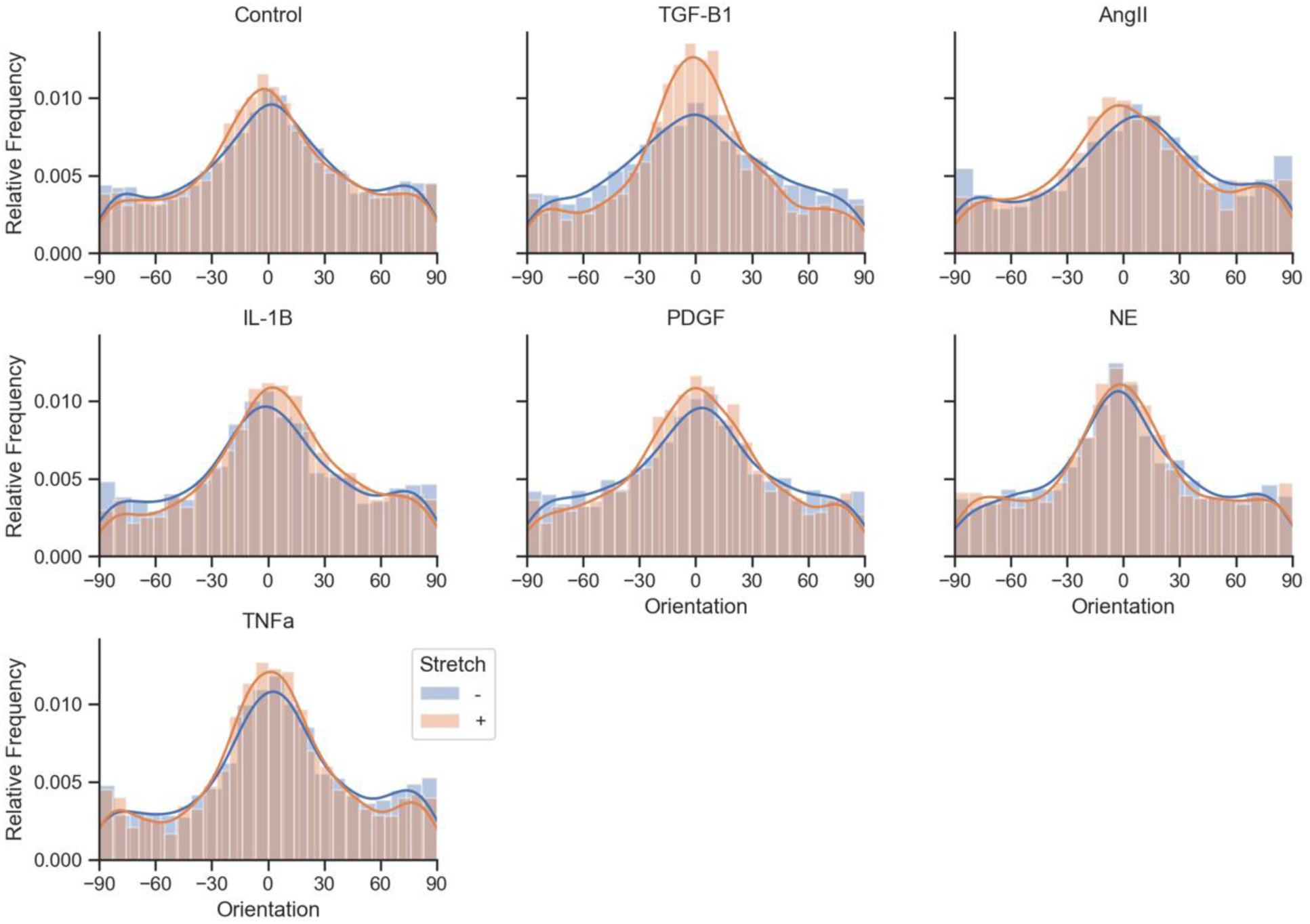
Fibroblasts maintain alignment along the axis of stretch in fibrin gels. Fluorescent images for fibroblast-seeded fibrin gels across all experimental groups were collected after conducting cell viability analysis for each sample (Fig. S2). The orientations of calcein-positive cells for one image per gel (400.2±183.9 cells per image, N=7 biological replicates) were calculated using CellProfiler software. Orientations are expressed as the angle in degrees from the axis of stretch, with 0° denoting cell alignment parallel to the direction of stretch, and ±90° denoting cell alignment perpendicular to the direction of stretch.

**Supplementary Figure S4.**
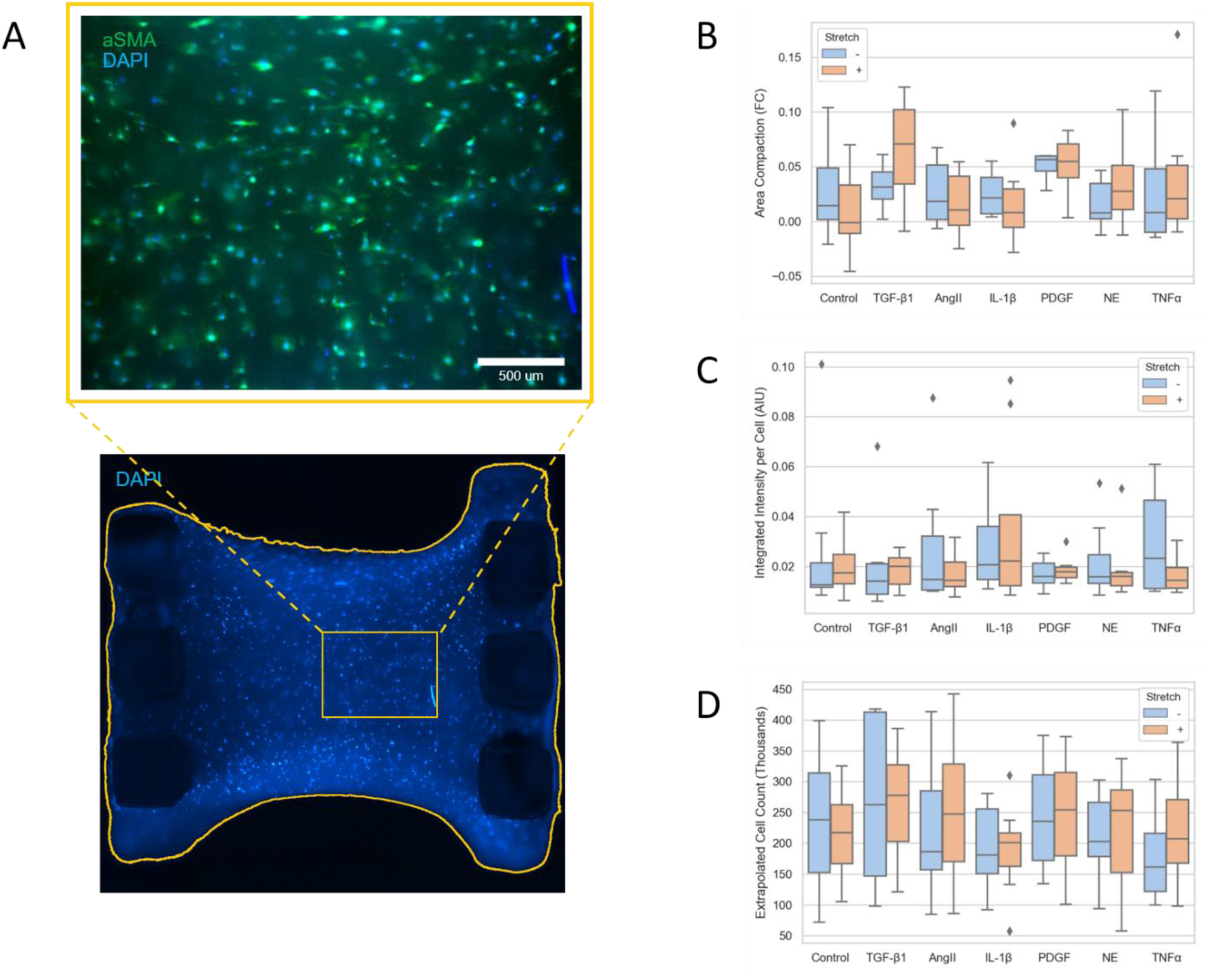
Immunocytochemistry and imaging analysis for fibroblast-seeded fibrin gels. Fibrin gels probed for αSMA and DAPI were analyzed for fluorescence intensity and cell count per image using CellProfiler software. Cell counts per image, as determined by DAPI-labeled nuclei, were extrapolated to measured areas of each gel in order to estimate total cell counts per gel. (B) Comparison of bulk gel compaction from before treatment to after treatment for high-dose agonist treatments and agonist-stretch combinations. (C) Comparison of fluorescence intensity for αSMA across high-dose agonist treatments and agonist-stretch combinations. Gel-averaged intensities were determined by averaging the integrated intensity of each cell across all cells in each image (N=8 biological replicates). (D) Comparison of extrapolated cell counts across high-dose agonist treatments and agonist-stretch combinations.

**Supplementary Figure S5.**
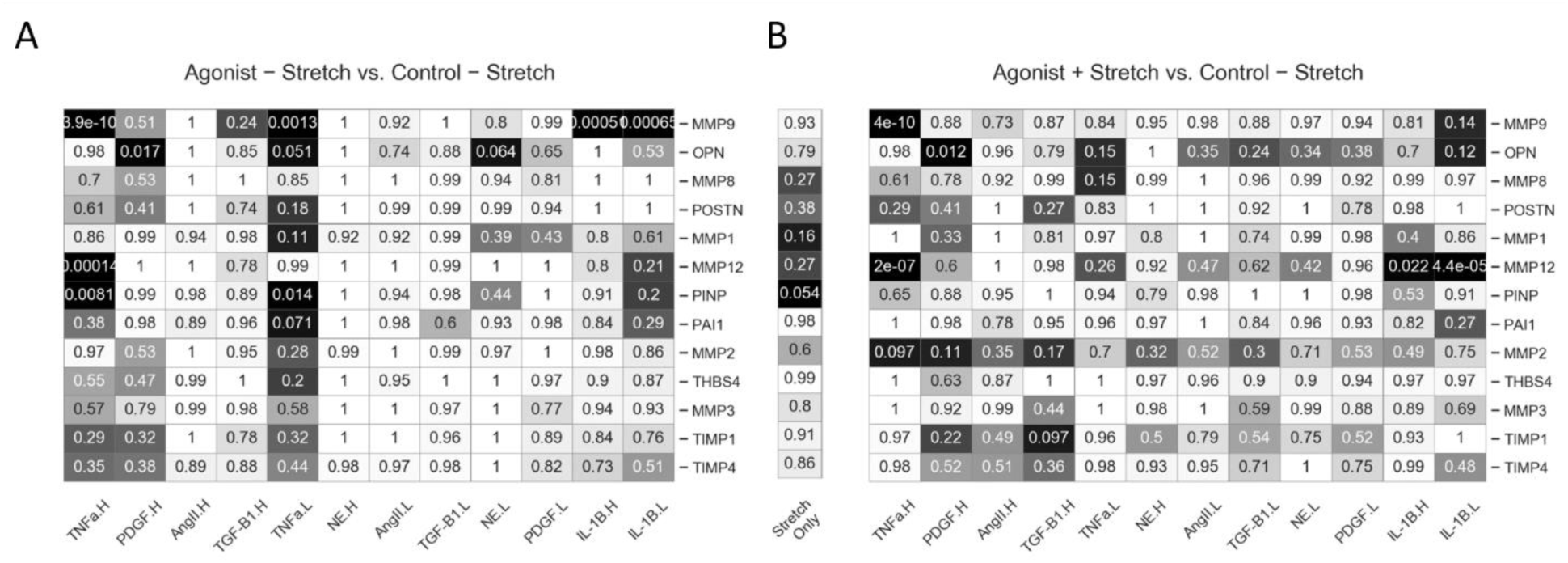
Statistical significance of individual agonists compared to un-stretched controls shown in Fig. 3. P-values of Dunnett’s test multiple comparisons are displayed for agonist treatments only versus un-stretched controls (A) and for agonist-stretch combinations versus un-stretched controls (B).

**Supplementary Figure S6.**
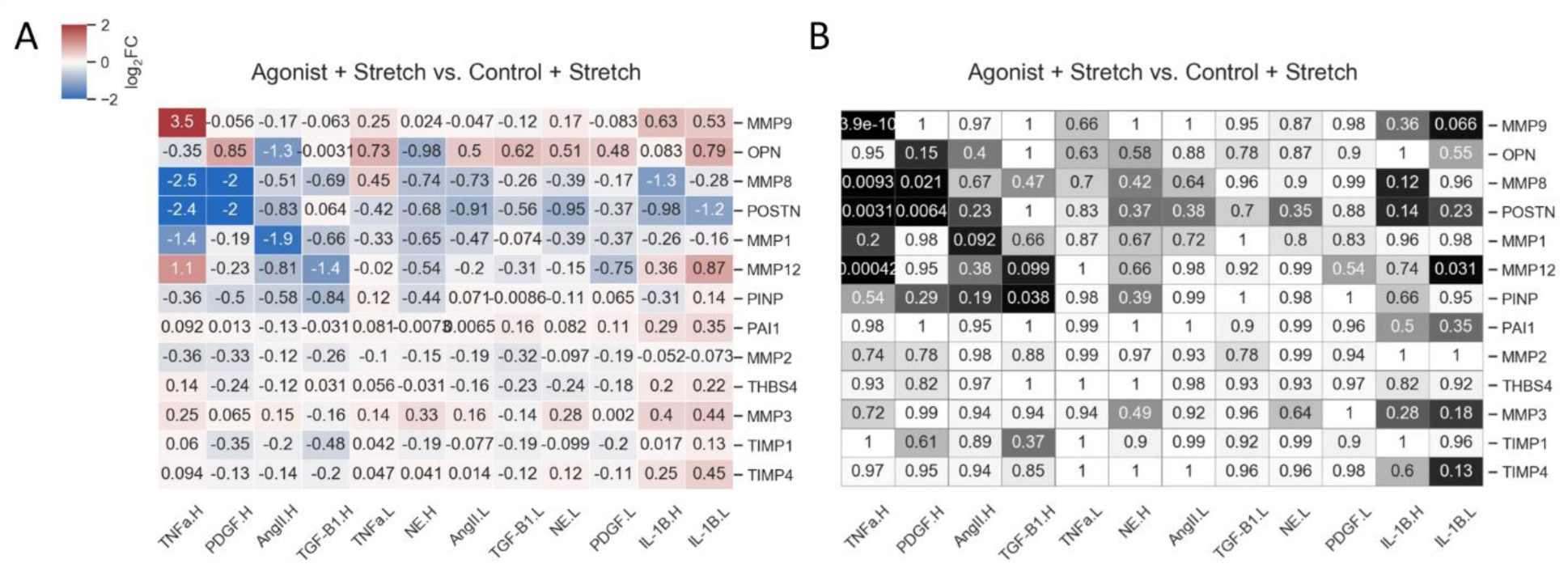
Effect and significance of agonist treatments on fibroblast secretomics in high-tension context. (A) Comparisons of agonist- and stretch-treated gels versus stretch-only gels across all secreted proteins, expressed as log_2_(fold-change). (B) P-values of contrasts for groups compared to stretched controls as determined by Dunnett’s tests.

## Notes

### Competing Interest Statement

The authors have declared no competing interest.

